# Dietary zinc protects against methotrexate-induced neurotoxicity in rats via modulation of oxidative stress, inflammation and neuronal energy metabolism

**DOI:** 10.1101/2023.04.24.538169

**Authors:** Adejoke Y. Onaolapo, Olufemi B Okunola, Anthony T. Olofinnade, Olakunle J. Onaolapo

## Abstract

**Background:** Increasing incidence of cancers and cancer chemotherapy-induced neurotoxicities makes it imperative to research compounds with neuroprotective potential that can do not impede therapy.

**Objective:** To examine the effect of dietary zinc supplementation on methotrexate-induced changes in neurobehaviour and neurochemistry and hippocampal morphology in rats.

**Methods:** Adult male rats were assigned into five groups of twelve animals each. Group were normal control and methotrexate control fed standard rodent chow and three groups of rats fed zinc supplemented diet at 25, 50 and 100 mg/kg of feed respectively. Standard and zinc supplemented diet were administered daily for 21 days. Animals in the normal control were administered intraperitoneal injection (i.p) of normal saline at 2ml/kg, while those in the methotrexate and zinc groups were administered i.p methotrexate at 20 mg/kg/day on days 19-21. On day 22, animals were exposed to behavioural paradigm (open field, Y-maze, radial arm maze, elevated plus maze and behavioural despair test). After the last behavioural test, animals were sacrificed and blood taken for the assessment of Tumour necrosis factor-α, interleukin 1β and interleukin 10), malondialdehyde levels and total antioxidant capacity. The hippocampus was either homogenised for the assessment of dopamine, serotonin, acetylcholine and brain derived neurotropic factor levels or processed for histological study.

**Results:** Dietary zinc protected against methotrexate-induced changes in weight, food intake, cognition, hippocampal histomorphology and neuron specific enolase immunohistochemistry.

**Conclusion:** Dietary zinc supplementation protects against methotrexate induced neurotoxicities by modulating oxidative stress, inflammatory markers brain neurotransmitters and metabolism in rats.

## 1.0 Introduction

Chemotherapy is a mainstay of cancer therapy, being used for the management of a variety of cancers and the prevention of tumor growth [1]. Most chemotherapeutic agents suppress the growth and proliferation of cancer cells; these mechanisms are however, not specific to cancer cells alone with non cancerous cells also impacted by the activity of these agents [2, 3]. The rising global prevalence of cancer means that more persons will need to be exposed to anticancer chemotherapy and by extension be at risk chemotherapy-induced toxicities. This chemotherapy related toxicities include nausea, vomiting, diarrhea, loss of appetite, alopecia and anaemia [4].

In the last few years there have also been reports suggesting an increase in the incidence of chemotherapy induced neurotoxicities [5–8]; while a number of these adverse neurological effects are first observed during therapy there have also been reports of chemotherapy induced neurotoxicity in cancer survivors [5–8]. In some instances it could necessitate the discontinuation of therapy [9]. Chemotherapy–induced neurotoxicity usually manifests as leucoencelopathy, cognitive deficits, peripheral neuropathy and mood disorders [9, 10]. Increased neuro-immune response, oxidative stress and the release of pro-inflammatory cytokines in the brain have been suggested as important mechanisms in the development of chemotherapy-induced neurotoxicity [9, 11–14]. Chemotherapeutic agents implicated in the development of chemotherapy induced neurotoxicity include doxorubicin, vinca alkaloids, taxanes, 5-fluorouracil, cyclophosphamide and methotrexate.

Methotrexate (MTX) is an antimetabolite used broadly in the management of cancers and also as an immunosuppressant in the managemet of rheumatoid arthritis [15]. It has been used alone or as an adjunct in the treatment of non-Hodgkin’s lymphoma, acute lymphoblastic leukemia breast, uterine, gestational and lung cancers [16]. In the last decade, there has been increasing evidence of methotrexate-related neurotoxicity [3, 15–18]. While age is a very important factor in the development of methotrexate-induced neurotoxicity, the severity of the neurological deficits observed with MTX has also been linked to dose, route of administration, and combination [19, 20]. Methotrexate-related neurotoxicity manifests a leucoencephalopathy that presents clinically as transient ischemic attacks, convulsion, encephalopathy, dementia, movement disorders. Its versatility in cancer chemotherapy and in the management of rheumatoid arthritis makes the search for neuroprotective agents that do not impede treatment imperative. The relationship amongst methotrexate induced neurotoxicity, oxidative stress, proinflammation and increased neuro-immune response has led to suggestions that compounds which have significant antiinflammatory, antioxidant and confirmed neuroprotective [15, 16, 20, 21] such as zinc could be efficient in preventing or mitigating the development of methotrexate induced neurotoxicity.

Zinc is a trace element essential for human survival, with its deficiency linked to the development of growth retardation, cognitive dysfunction and mood disorders [22]. Zinc has been linked with the synthesis and normal activity of a variety of enzymes, playing crucial roles in the suppression of inflammation, oxidative stress, hyperactive immune system, and in the maintenance of glucose and energy metabolism [22, 23]. In the brain, zinc is a structural component of a protein and contributes significantly to the normal functioning of transcription factors and enzymes [24–26]. It has also been shown to be present in the brain at higher concentrations compared to its concentration in serum. The amygdala and hippocampus have the highest concentration of zinc while the brain’s extracellular fluid has the lowest concentration [24–26]. The neuroprotective effects of zinc have also been reported severally [27–30]. While there have been reports of the ameliorative potential of zinc in methotrexate induced intestinal damage [31, 32]. There is a dearth of information on its possible effect in protecting against methotrexate induced neurotoxicity, hence this study.

## 2.0 Materials and Methods

### 2.1 Drugs and Reagents

Zinc (as ZN gluconate caplet, 100 mg, Masson Vitamins Inc., Florida, USA), containing 14 mg of elemental zinc/100 mg of zinc gluconate.Methotrexate injection 50 mg/ 2 ml, Aspar Pharmaceuticals, Gujarat, India. Assay kits for lipid peroxidation (malondialdehyde), interleukin-10, tumour necrosis factor alpha and total antioxidant capacity (Biovision Inc., Milpitas, CA, USA).

### 2.2 Animals

Male Wistar rats were obtained from Empire Breeders, Orioke,-Ara Osun State and housed in metal cages at room temperature (25.°C ±2.5°C) with 12 hours of light. Rats had free access to food and tap water. All procedures would be in adherence to approved protocols of the Ladoke Akintola University of Technology, and also as prescribed within the European Council Directive (EU2010/63) for the use and care of laboratory animals.

### 2.3 Diet

Diet consisted of commercially available standard rodent chow (Top Feeds®, Nigeria). During the experimental period, animals were fed standard diet (29% protein, 11% fat, 58% carbohydrate, and 10 mg of zinc gluconate/kg of feed) or zinc-supplemented diet at 25, 50 and 100 mg of Zinc gluconate/kg of feed and corresponding to 3.5, 7 and 14 mg of elemental zinc supplementation. Feed was administered ad libitum. The concentration of ZN in the diet was patterned after the study by Onaolapo et al. [33].

### 2.4 Experimental methodology

Sixty young adult male rats weighing 120—150 g each were randomly assigned into five groups of twelve (n=12) animals each. Group were normal control and methotrexate control fed standard rodent chow and three groups of rats fed zinc supplemented diet at 25, 50 and 100 mg/kg of feed respectively. Standard and zinc supplemented diet were administered daily for 21 days. Animals in the normal control were administered intraperitoneal injection (i.p) of normal saline at 2ml/kg, while those in the methotrexate and zinc groups were administered i.p methotrexate at 20 mg/kg/day on days 19-21. At the end of the experimental period, animals were exposed to varied behavioural paradigm (open field, Y-maze, radial arm maze and elevated plus maze). Twenty-four hours after the last behavioural tests, animals were sacrificed by cervical dislocation and blood taken through an intracardiac puncture for the assessment of levels of inflammatory markers (Tumour necrosis factor-α, interleukin 1β and interleukin-10), lipid peroxidation (measured as malondialdehyde levels) and antioxidant capacity (measured as total antioxidant capacity). The brain was removed, observed grossly, and weighed. Sections of the hippocampus were processed for paraffin-embedding, cut at 5 µm and stained for histological study. Supernatant from homogenates of the hippocampus was used to assess the activity of brain neurotransmitters (dopamine, serotonin, acetylcholine and brain derived neurotropic factor).

### 2.5 Body weight and food intake Determination

Body weight was measured weekly while food intake was measured daily using an electronic weighing balance as described previously [34–37]. Calculated for each rat was their relative change in body weight or food intake using the equation below, following which results for all animals were computed to determine the statistical mean.

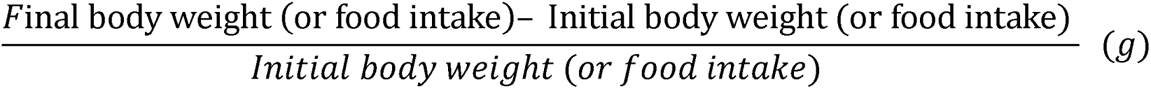

### 2.6 Behavioural tests

Behavioural tests were carried out in this sequence: 1) elevated plus maze, 2.) open field and 3) memory tests, 4) behavioural despair.

#### 2.6.1 Anxiety Model: Elevated plus-maze

The elevated plus-maze (EPM) is a plus-shaped paradigm with four arms arranged at right angles to each other. The EPM is used to measure anxiogenic or anxiolytic behaviours which are scored as time spent in the closed or open arms within a 5 minute period of exposure to the maze. Anxiety behaviours were scored as previously described [38–41].

#### 2.6.2 Open field behaviours

Exposure to the Open-field paradigm in rodent is used to measure central behaviours. Self-grooming is used to depict stereotypic behaviours. Generally central behaviours are indicative of a rat’s ability to explore open spaces. Ten minutes of open-field exposure was used to assess and score horizontal locomotion, rearing and self-grooming behaviours. The open-field paradigm is a rectangular box with a white painted hard floor that measuring 72 x 72 x 26 cm. The hard wood floor was divided by permanent red markings into 16 equal squares. The placement of animal within the box, its movement patterns and scoring are as described [42–44].

#### 2.6.3 Memory tests (Y-and Radial arm- maze)

The Y-maze and radial arm mazes are employed for the measurement of spatial working memory. Spatial working-memory is assessed by monitoring spontaneous alternation behaviour of a rat placed in the maze for five minutes. Spontaneous alternation behaviour assesses the propensity of rodents to alternate conventionally non-reinforced choices of the Y or radial arm maze on successive chances. The sequence of arm entries is assessed and scored as previously described [41, 42, 45].

The radial arm maze has eight equidistantly-spaced arms having a length of at least 3 cm long. Each arm radiating from a central platform. Each rat is placed on the central platform and allowed free movement in respective arms during which its behaviours are recorded. Working memory is scored when the rat enters each arm once as described previously [45, 46].

#### 2.6.4 Tail Suspension test

\The tail suspension test measures behavioural despair. Using previously described protocols [36], which involved suspending animals by fastening them securely to a flat platform by the tip of their tail for 6 minutes. Total time spent immobile was then measured during the 6-minute period of the testing session

### 2.7 Biochemical Test

#### 2.7.1 Estimation of MDA content (Lipid peroxidation)

Lipid peroxidation level were measured as malondialdehyde content as described previously [47–49].

#### 2.7.2 Antioxidant activity

Total antioxidant capacity was measured with commercially available assay kit. Colour changes were monitored as described previously [49–51].

#### 2.7.3 Acetylcholine, dopamine, serotonin and BDNF levels

Supernatants decanted from homogenate of the hippocampus was used to assay for levels of acetylcholine, dopamine, serotonin and brain derived neurotrophic factor using commercially available Enzyme linked immunosorbent assay kits according to the instructions of the manufacturer (ABCAM, Cambridge UK)

#### 2.7.4 Tumour necrosis factor-α, Interleukin (IL) –10 and Interleukin 1β

Tumour necrosis factor-α, interleukin (IL)-10 and Interleukin1 β level were measured using enzyme-linked immunosorbent assay (ELISA) techniques with commercially available kits (Enzo Life Sciences Inc. NY, USA) designed to measure the ‘total’ (bound and unbound) amount of the respective cytokines as previously described [52, 53].

#### 2.7.5 Protocol for neuron specific enolase (NSE) Immunohistochemistry

Neuron specific enolase (NSE) immunohistochemistry was carried out using NSE primary monoclonal antibody and the Novocastra™ and Novolink DM polymer detection system (Leica Biosystems, UK) as described previously [54, 55].

### 2.8 Photomicrography

Sections of the hippocampus processed for histology were examined using a Sellon-Olympus trinocular light microscope (XSZ-107E, China) with a digital camera (Canon Powershot 2500) attached. Histopathological changes were assessed by an observer blinded to the groupings.

### 2.9 Statistical analysis

Data were analysed with Chris Rorden’s ANOVA for windows (version 0.98). Data analysis was by One-way analysis of variance (ANOVA) and post-hoc test (Tukey HSD) was used for within and between group comparisons. Results were expressed as mean ± S.E.M. and p < 0.05 was taken as the accepted level of significant difference from control.

## 3.0 Results

### 3.1 Effects of Zinc supplementation on body weight and food intake

Figure 1 shows the effect dietary zinc supplementation on relative change in body weight (upper panel) and change in food intake (lower panel) in methotrexate (MTX)-treated rats. There was a significant [F (4, 45) = 163, p < 0.001] decrease in weight gain with MTX, MTX/ZN 25, MTX/ZN50 and MTX/ZN100 compared to control. Compared to MTX, body weight increased significantly with at MTX/ZN50 and MTX/ZN100. Food intake decreased significantly [F (4, 45) = 171, p < 0.001] MTX, and MTX/ZN at 25 mg/kg and an increase with MTX/ZN50 and MTX/ZN100 compared to control. Compared to MTX, food intake increased significantly at MTX/ZN25, MTX/ZN50 and MTX/ZN100.

**Figure 1:**
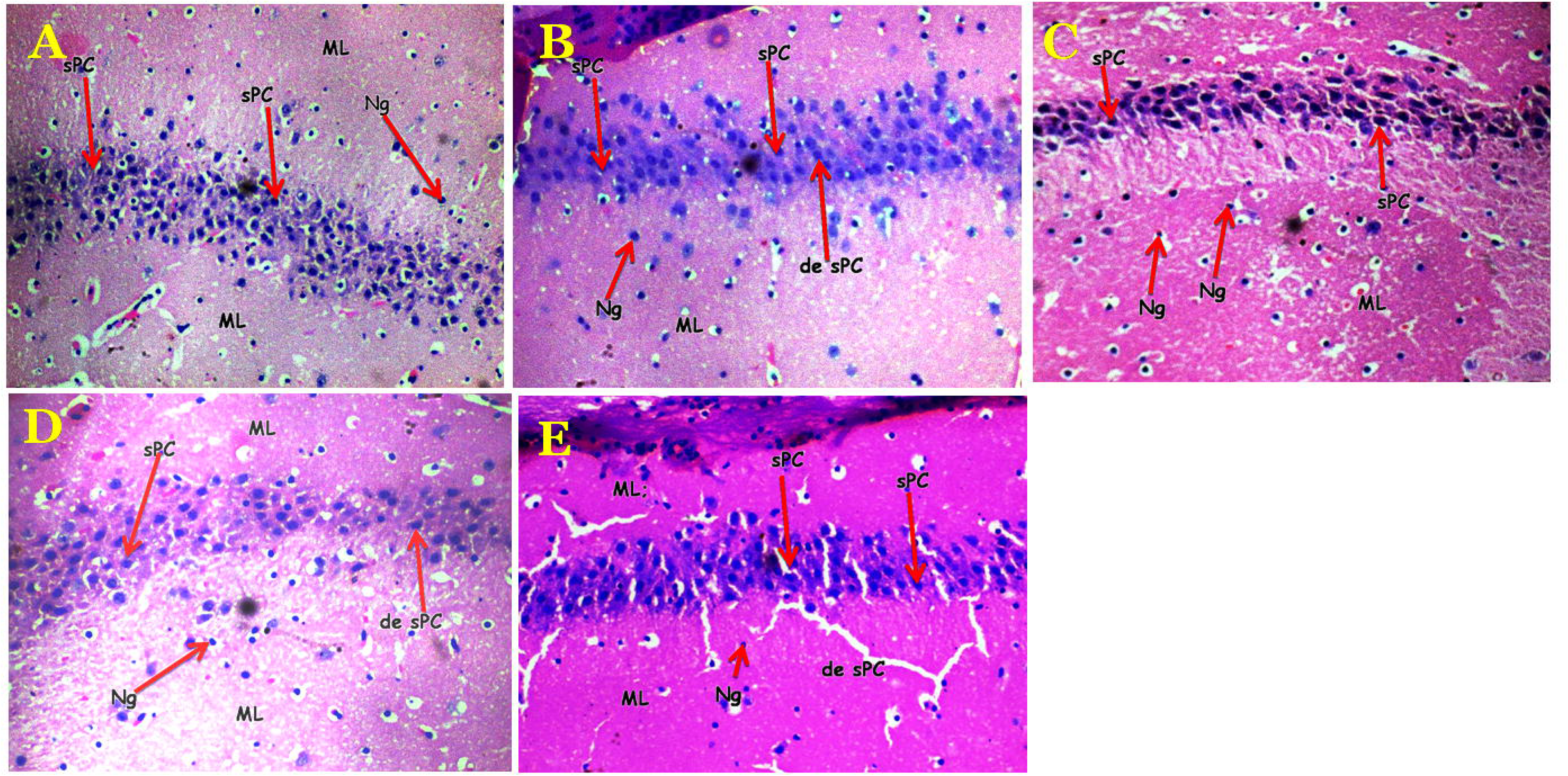
effect dietary zinc supplementation on relative change in body weight (upper panel) and change in food intake (lower panel) in methotrexate (MTX)-treated rats. Each bar represents Mean ± S.E.M, *p < 0.05 significant difference from control, #p<0.05 significant difference from MTX, number of mice per treatment group =10. MTX: Methotrexate.

### 3.2 Effect of zinc on locomotion in methotrexate-treated rats

Figure 2 shows the effect dietary zinc supplementation on horizontal locomotion measured as number of line crossings (upper panel)and vertical locomotion (rearing) in MTX-treated rats. There was a significant [F (4, 45) = 18.3, p < 0.001] decrease in line crossing with MTX, MTX/ZN25, MTX/ZN50 and MTX/ZN100 compared to control. Compared to MTX, line crossing increased significantly with MTX/ZN 25 and decreased with MTX/ZN50 and MTX/ZN100. Vertical locomotion (rearing) decreased significantly [F (4, 45) = 75.7, p < 0.001] with MTX, MTX/ZN25, MTX/ZN50 and MTX/ZN100 mg/kg of feed compared to control. Compared to MTX, rearing activity increased significantly at MTX/ZN25 and decreased with MTX/ZN50 and MTX/ZN100.

**Figure 2:**
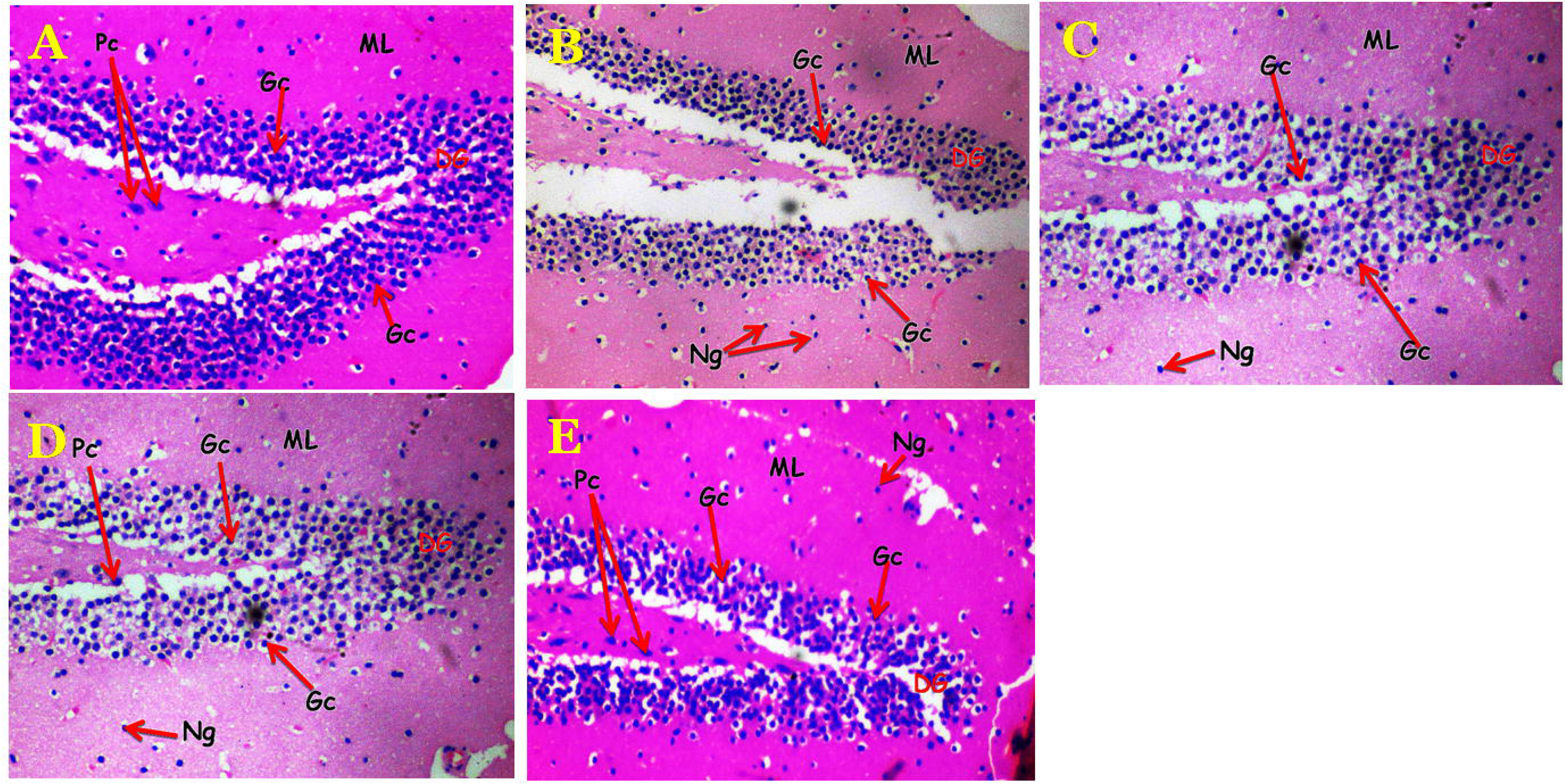
Effect of zinc (ZN) on horizontal (upper panel) and vertical (Lower panel) locomotion in MTX-treated rats. Each bar represents Mean ± S.E.M, *p < 0.05 significant difference from control, #p<0.05 significant difference from MTX, number of mice per treatment group =10. MTX: Methotrexate.

### 3.3 Effect of zinc on self-grooming and behavioural despair

Figure 3 shows the effect dietary zinc supplementation on self-grooming behaviours (upper panel) and immobility time in the tail suspension test (lower panel) in methotrexate treated rats. There was a significant [F(4,45) = 19.7, p < 0.001] decrease in self-grooming with MTX, MTX/ZN 25, MTX/ZN 50 and MTX/ZN 100 compared to control. Compared to MTX, self-grooming increased significantly at MTX/ZN 25, MTX/ZN 50 and MTX/ZN 100.

**Figure 3:**
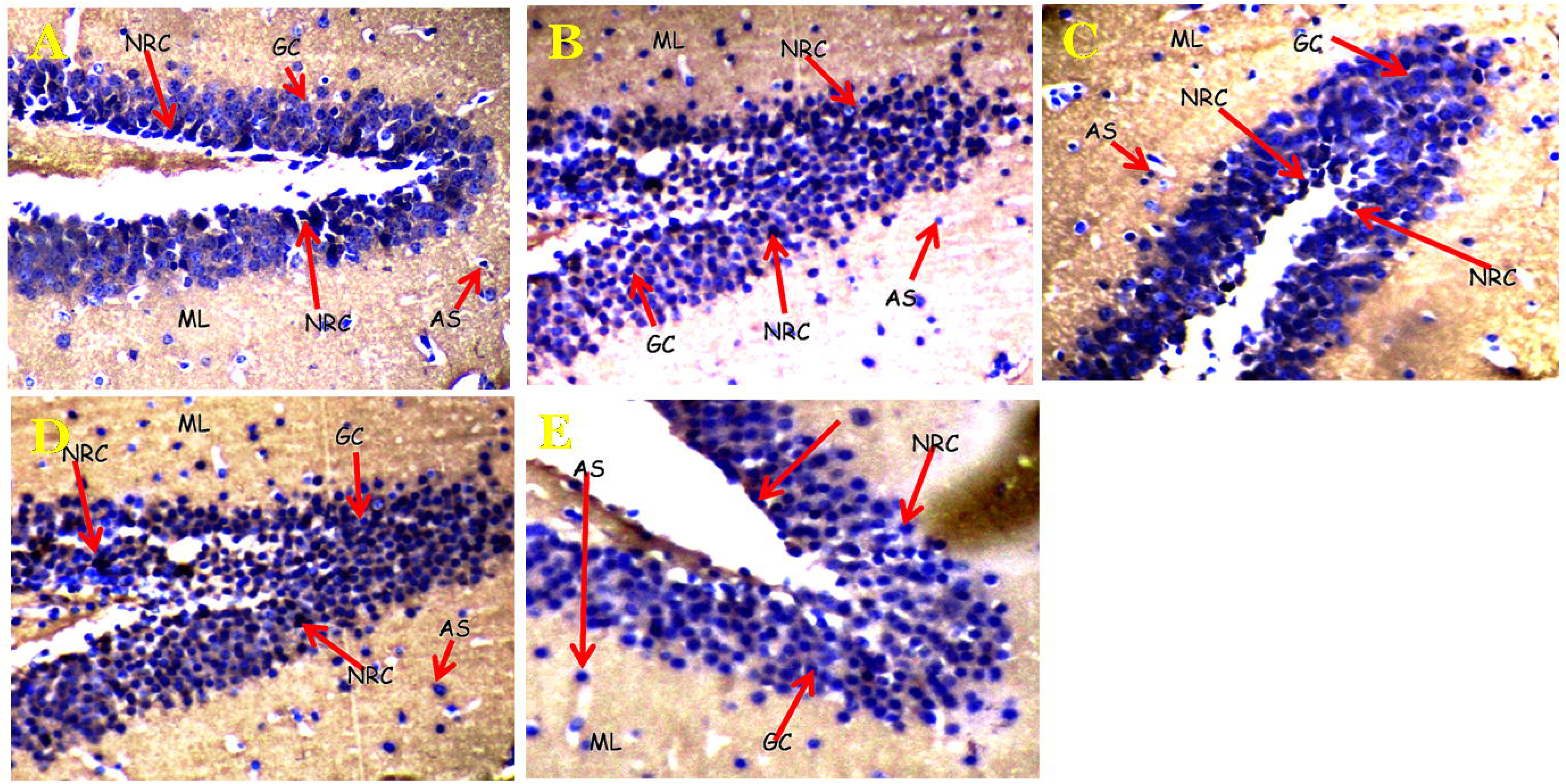
Effect of zinc (ZN) on self-grooming (upper panel) and immobility time in the tail suspension test (Lower panel) in MTX-treated rats. Each bar represents Mean ± S.E.M, *p < 0.05 significant difference from control, #p<0.05 significant difference from MTX, number of mice per treatment group =10. MTX: Methotrexate.

Immobility time increased significantly [F(4,45) = 126.32, p < 0.001] with MTX, MTX/ZN 25, and MTX/ZN 50 compared to control. Compared to MTX, immobility time decreased significantly at \MTX/ZN 25, MTX/ZN 50 and MTX/ZN 100.

### 3.4 Effect of zinc supplementation on spatial-working memory

Figure 4 shows the effect dietary zinc supplementation on spatial working memory in the Y (upper panel) and radial arm (lower panel) maze in methotrexate treated rats. There was a significant [F(4,45) = 102, p < 0.001] decrease in spatial working memory with MTX, MTX/ZN 50 and MTX/ZN 100 and an increase with MTX/ZN 25 compared to control. Compared to MTX, spatial working memory in the Y maze increased significantly with MTX, MTX/ZN25 MTX/ZN50 and MTX/ZN100.

**Figure 4:**
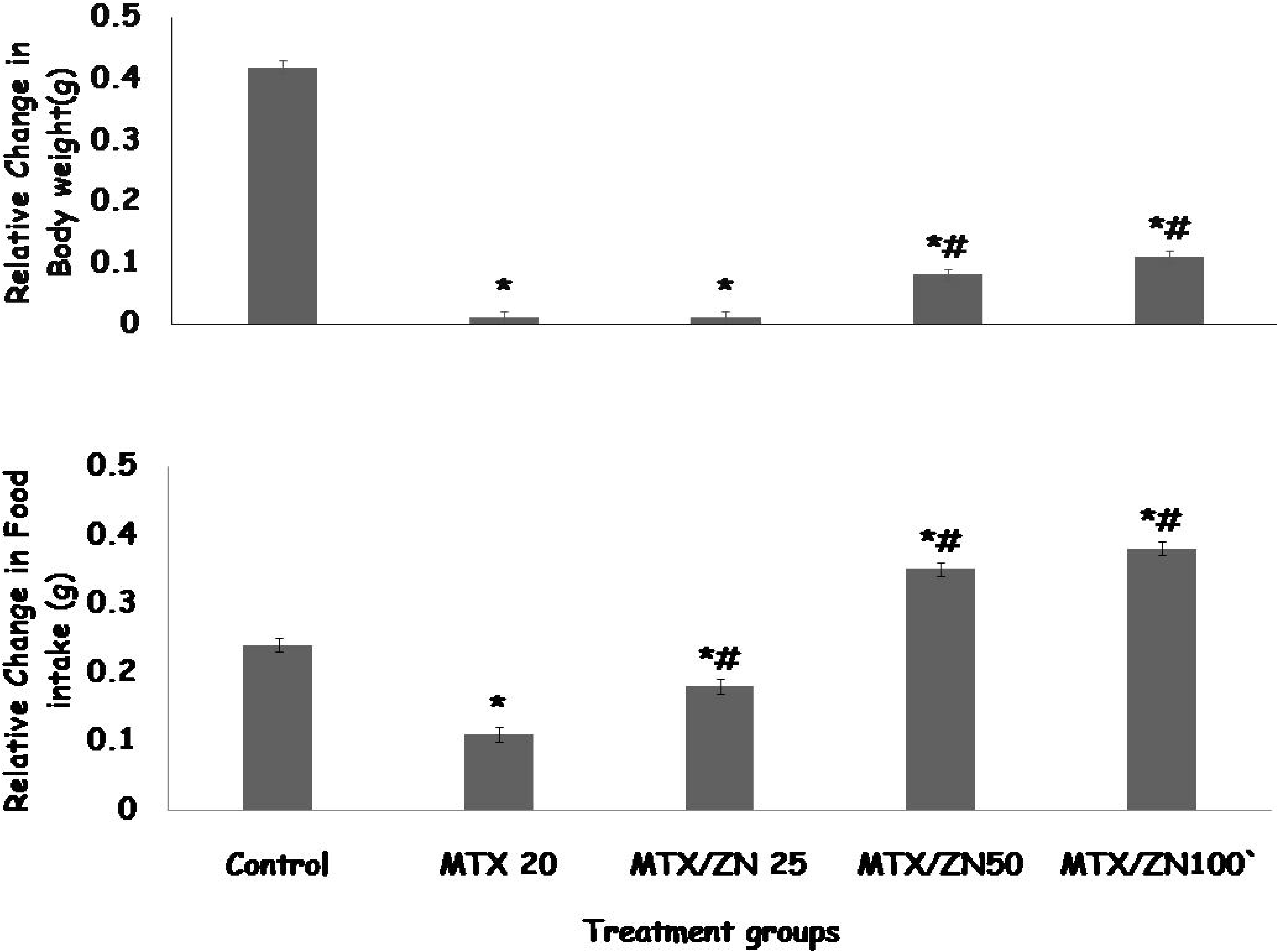
Effect of zinc on spatial working memory in the Y-maze (upper panel) and radial arm maze (lower panel) in methotrexate treated rats. Each bar represents Mean ± S.E.M, *p < 0.05 significant difference from control, #p<0.05 significant difference from MTX number of mice per treatment group =10. MTX: Methotrexate.

Spatial working memory in the radial arm maze decreased significantly [F(4,45) = 32.4, p < 0.001] with MTX, MTX/ZN 50 and MTX/ZN 100 and increased with MTX/ZN 25 compared to control. Compared to MTX, spatial working memory in the radial arm maze increased significantly with MTX, MTX/ZN25 MTX/ZN50 and MTX/ZN100.

### 4.7 Effect of zinc on time spent in the arms of the elevated plus-maze

Figure 5 shows the effect dietary zinc supplementation on time spent in the open arm (upper panel) and closed arm of the elevated plus maze in methotrexate-treated rats. There was a significant [F(4,45) = 25.7, p < 0.001] decrease in time spent in the open arm with MTX, MTX/ZN 50 and 1MTX/ZN 100 and an increase with MTX/ZN25 compared to control. Compared to MTX, open arm time increased significantly with MTX/ZN25, MTX/ZN50 and MTX/ZN100.

**Figure 5:**
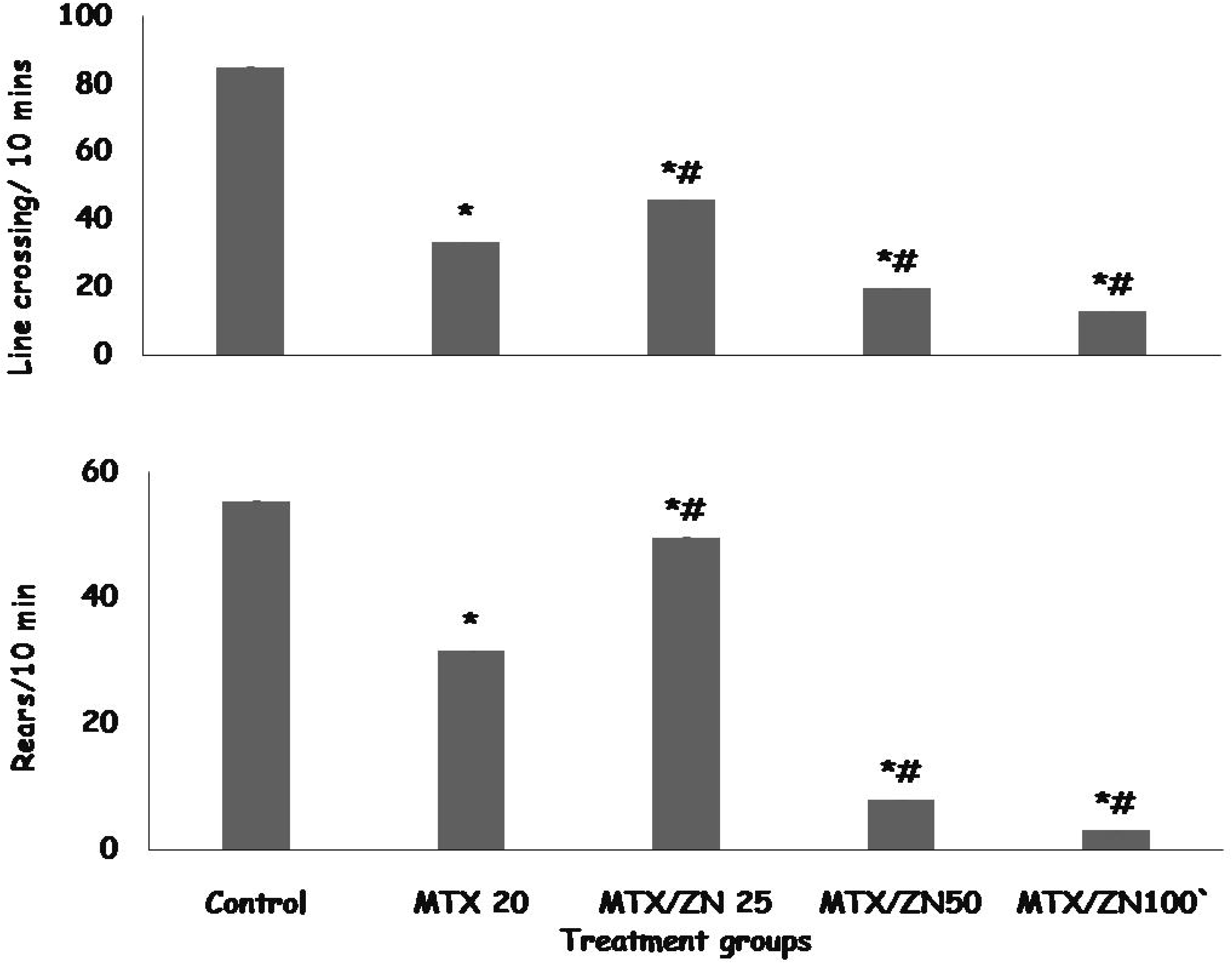
Effect of zinc on time spent in the open arm (upper panel) and closed arm (lower panel) of the elevated plus maze in methotrexate-treated rats. Each bar represents Mean ± S.E.M, *p < 0.05 significant difference from control, #p<0.05 significant difference from MTX. number of mice per treatment group =10. MTX: Methotrexate.

Closed arm time increased significantly [F(4,45) = 22.1, p < 0.001] with MTX, MTX/ZN50 and MTX/ZN100 compared to control. Compared to MTX, closed arm time decreased significantly with MTX/ZN25.

### 4.6 Effect of zinc on biochemical assays

Table 1 shows the effect dietary zinc supplementation on lipid peroxidation (malondialdehyde), total antioxidant capacity, interleukin-10 and tumour necrosis factor-α levels in methotrexate (MTX)-treated rats. There was a significant [F(4,45) = 12.1, p < 0.001] increase in malondialdehyde levels with MTX, and MTX/ZN50 and MTX/ZN100 and a decrease with MTX/ZN25 compared to control. Compared to MTX, malondialdehyde levels decreased significantly with MTX/ZN25, MTX/ZN50 and MTX/ZN100.

**Table 1:** Effect of zinc on Biochemical assays

| Groups | MDA (uM) | TAC(MmTE) | TNF-α (ng/L) | IL-10 (pg/ml) | IL-1β (pg/ml) |
| --- | --- | --- | --- | --- | --- |
| Control | 5.59±1.10 | 5.57±0.23 | 55.48±1.67 | 68.62±2.76 | 30.04±1.22 |
| MTX 20 | 15.30±1.10* | 1.80±0.14* | 80.23±1.22* | 32.23±1.30* | 53.22±1.76* |
| MTX/ZN 25 | 4.10±0.35*# | 22.19±0.17*# | 50.25±1.11*# | 176.44±1.46* | 26.32±0.75*# |
| MTX/ZN50 | 7.40±0.30*# | 4.22±0.12*# | 55.19±1.20* | 43.63±1.43*# | 12.11±1.23* |
| MTX/ZN100 | 9.90±0.20# | 3.31±0.10# | 54.74±2.25# | 40.21±1.34*# | 10.03±1.11* |

Data presented as Mean ± S.E.M, \* $p < 0.05$ vs. control, # $p < 0.05$ vs. MTX, Number of mice per treatment group=10, MDA: malondialdehyde, TAC; total antioxidant capacity, IL-10: Interleukin -10, TNF-α ; Tumour necrosis factor-alpha, TE: trollox equivalent.

Total antioxidant capacity (TAC) decreased significantly [F(4,45) = 16.9, p < 0.001] with MTX, and MTX/ZN50 and MTX/ZN100 and increased with MTX/ZN25 compared to control. Compared to MTX, TAC levels increased significantly with MTX/ZN25, MTX/ZN50 and MTX/ZN100.

Tumour necrosis factor--α (TNF-α) increased significantly [F (4,45) = 43.1, p < 0.001] with MTX, MTX/ZN50 and MTX/ZN100 compared to control. Compared to MTX, TNF-αlevels decreased significantly with MTX/ZN25.

Interleukin 10 (IL-10) levels decreased significantly [F(4,45) = 16.21, p < 0.001] with MTX, MTX/ZN50 and MTX/ZN100 and increased with MTX/ZN25 compared to control. Compared to MTX, IL-10 levels increased with MTX/ZN25, MTX/ZN50 and MTX/ZN100.

Interleukin 1 beta (IL-1β) levels increased significantly [F (4,45) = 23.22, p < 0.001] with MTX, MTX/ZN50 and MTX/ZN100 and decreased with MTX/ZN25 compared to control. Compared to MTX, IL-1β levels decreased significantly with MTX/ZN25.

### 4.7 Effect of zinc supplementation on neurotransmitter levels

Table 2 shows the effect zinc supplementation on neurotransmitter levels in the hippocampus in methotrexate treated rats. There was a significant increase in acetylcholine levels with MTX/ZN25 and a decrease with MTX, MTX/ZN50 and MTX/100 compared to control. Compared to MTX, acetylcholine levels increased with MTX/ZN25, MTX/ZN50 and MTX/ZN 100.

**Table 2:** Effect of dietary zinc supplementation on neurotransmitter levels in the hippocampus

| Groups | Acetylcholine<br>(pmol/mg<br>tissue) | Dopamine<br>(pmol/mg tissue) | BDNF<br>(pmol/mg<br>tissue) | Serotonin<br>(pmol/mg<br>tissue) |
| --- | --- | --- | --- | --- |
| Control | 3.52±0.11 | 40.02±1.90 | 23.11±0.10 | 6.33±0.10 |
| MTX | 0.54±0.10* | 22.12±0.25* | 12.21±0.20* | 3.22±1.10* |
| MTX/Zn25 | 5.22±0.12*# | 30.22±0.14*# | 25.22±0.30*# | 8.16±2.06# |
| MTX/Zn50 | 2.33±0.11*# | 27.00±0.16*# | 16.10±0.24*# | 5.40±1.04# |
| MTX/Zn100 | 1.20±0.11*# | 25.55±0.13*# | 14.12±0.25*# | 4.49±1.01# |

Data presented as Mean ± S.E.M, \* $p < 0.05$ vs. control, # $p < 0.05$ vs. MTX, Number of mice per treatment group=10, ZN: Zinc, BDNF: Brain derived neurotropic factor.

Dopamine levels decreased with MTX, MTX/ZN25, MTX/ZN50 and MTX/ZN100 compared to control. Compared to MTX, dopamine levels increased with MTX/ZN25, MTX/ZN50 and MTX/ZN100.

Brain derived neurotropic factor (BDNF) levels decreased with MTX, MTX/ZN25, MTX/ZN50 and MTX/ZN100 compared to control. Compared to MTX, BDNF levels increased with MTX/ZN25, MTX/ZN50 and MTX/ZN100.

Serotonin levels were significantly decreased with MTX, compared to control. Compared to MTX, serotonin levels increased with MTX/ZN25, MTX/ZN50 and MTX/ZN100.

### 4.8 Effect of zinc supplementation on hippocampal histomorphology

Figures 6(a-e) and 7(a-e) show representative photomicrographs of haematoxylin and eosin stained sections of the dentate gyrus and cornus ammonis 3 regions of the rat hippocampus. Examination of the slides of animals in the control group (6a, 7a) revealed the line of pyramidal neurons that predominate the cornus ammonis 2 region (Figure 6) as well as the triangular shaped layer of small granule cell neurons that make up the dentate gyrus (Figure 7). Also observed are glia neurons and neuronal processes in the molecular layer that lie between the compact zones of the cornus ammonis and dentate gyrus. In the groups administered methotrexate (6b, 7b) a graded loss of neurons and neuronal degeneration is observed. This is evidenced by poorly-staining nucleoli, loss of oval to round shape of the granule neuron, shrunken nucleus, absence of the close-knit arrangement of the granule neurons. In the cornus ammonis 2 region there was loss of pyramidal neurons with remaining pyramidal neurons at varying degrees of degeneration. In the groups fed zinc supplemented diet an amelioration of methotrexate-induced changes in the cornus ammonis 2 (Figure 6c–e) and dentate gyrus (Figure 7c–e) regions were observed.

**Figure 6(A-E):**
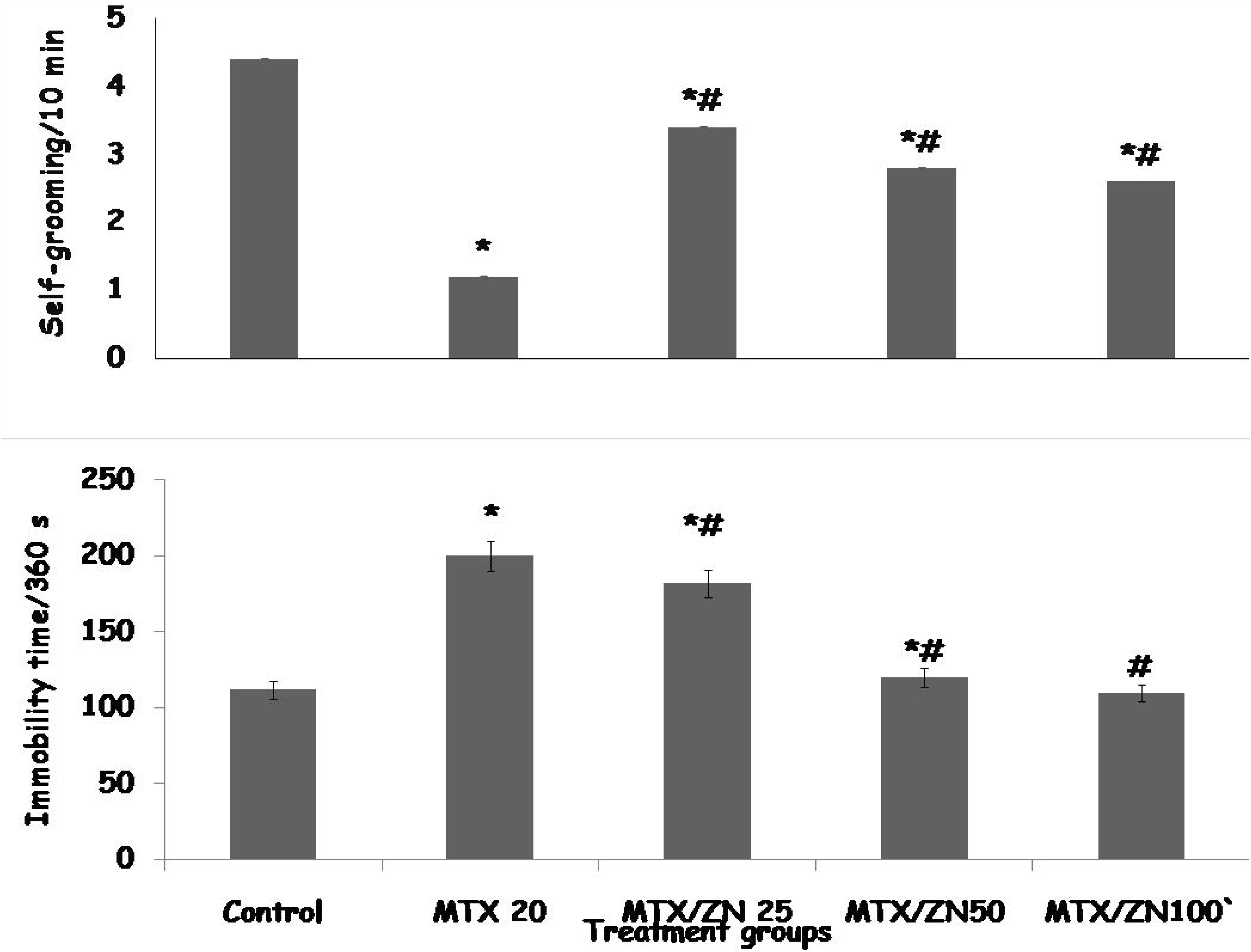
Histomorphology of the dentate gyrus of the hippocampus. A: Control, B: Methotrexate Control. C: MTX/ZN25. D: MTX/ZN50, E: MTX/ZN100. Photomicrograph showing pyramidal cells (Pc), small granule cells (Gc) within the dentate gyrus proper and neuroglia (Ng) scattered within the molecular layer (ML). MTX: Methotrexate. ZN: Zinc. Haematoxylin and Eosin staining method, Magnification=x160

**Figure 7(A-E):**
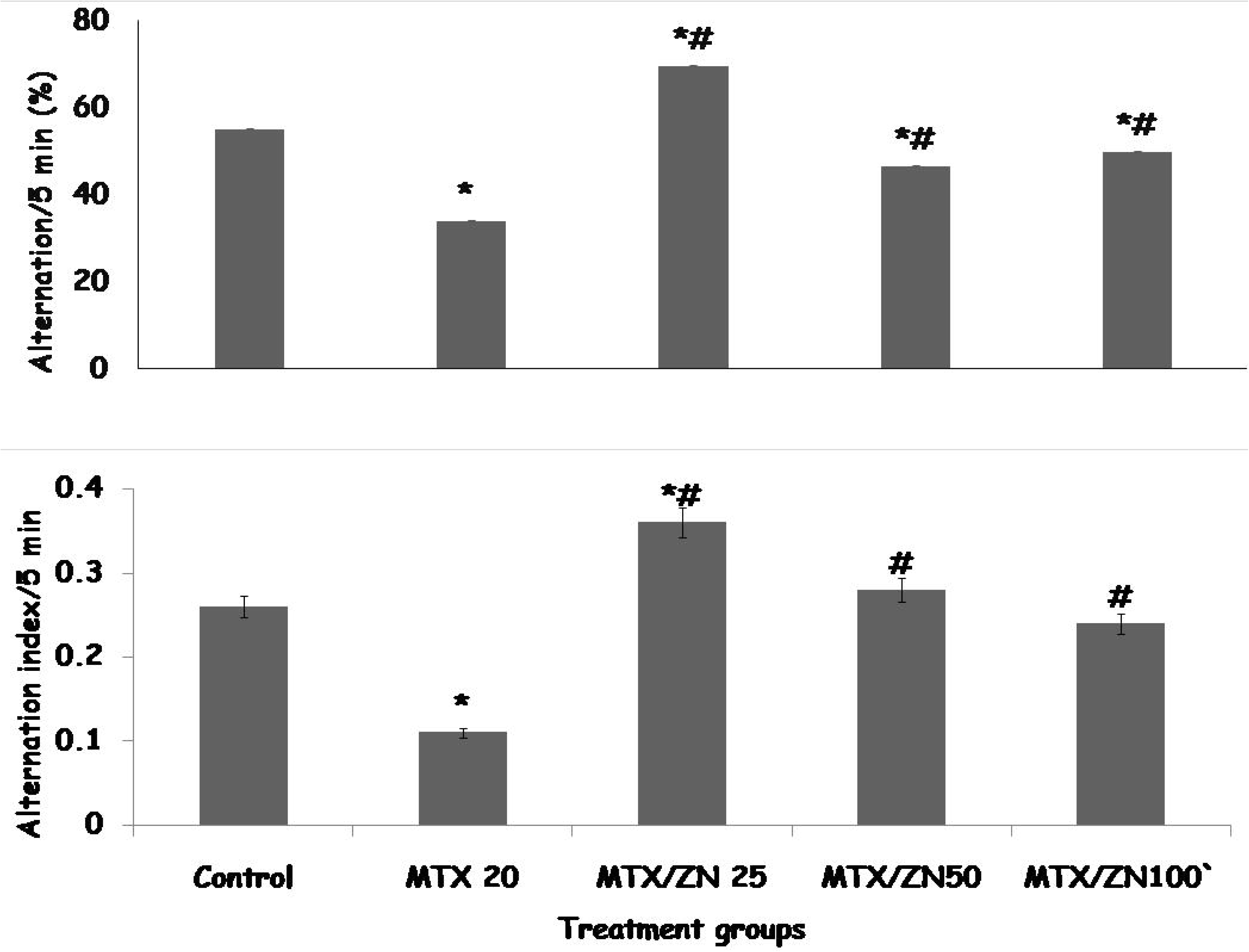
Histomorphology of the cornus ammonis 3 region of the hippocampus. A: Control, B: Methotrexate Control. C: MTX/ZN25. D: MTX/ZN50, E: MTX/ZN100. Photomicrograph showing small pyramidal cells (sPc), and neuroglia scattered within the molecular layer (ML). MTX: Methotrexate. ZN: Zinc. Haematoxylin and Eosin staining method, Magnification=x160

### 4.9 Effect of zinc supplementation on Neuron Specific Enolase (NSE) immunoreactivity in the hippocampus

Examination of the NSE-stained sections of the hippocampus (Figure 8a-e) of animals in the control group (figure 8a) revealed presence of NSE reactive neurons with granule cell morphologies (medium-sized round to oval cell bodies with brown-staining cytoplasm and blue staining nucleus) in the dentate gyrus, in the methotrexate control (figure 8b) there was an increase in NSE-reactivity with a cells in the dentate gyrus showing brown to black cytoplasm and pale blue staining nucleus. In the groups fed zinc supplemented diet (figure 8c-e) a reversal of NSE reactivity in the dentate gyrus to levels observed in the control group were observed.

**Figure 8(A-E):**
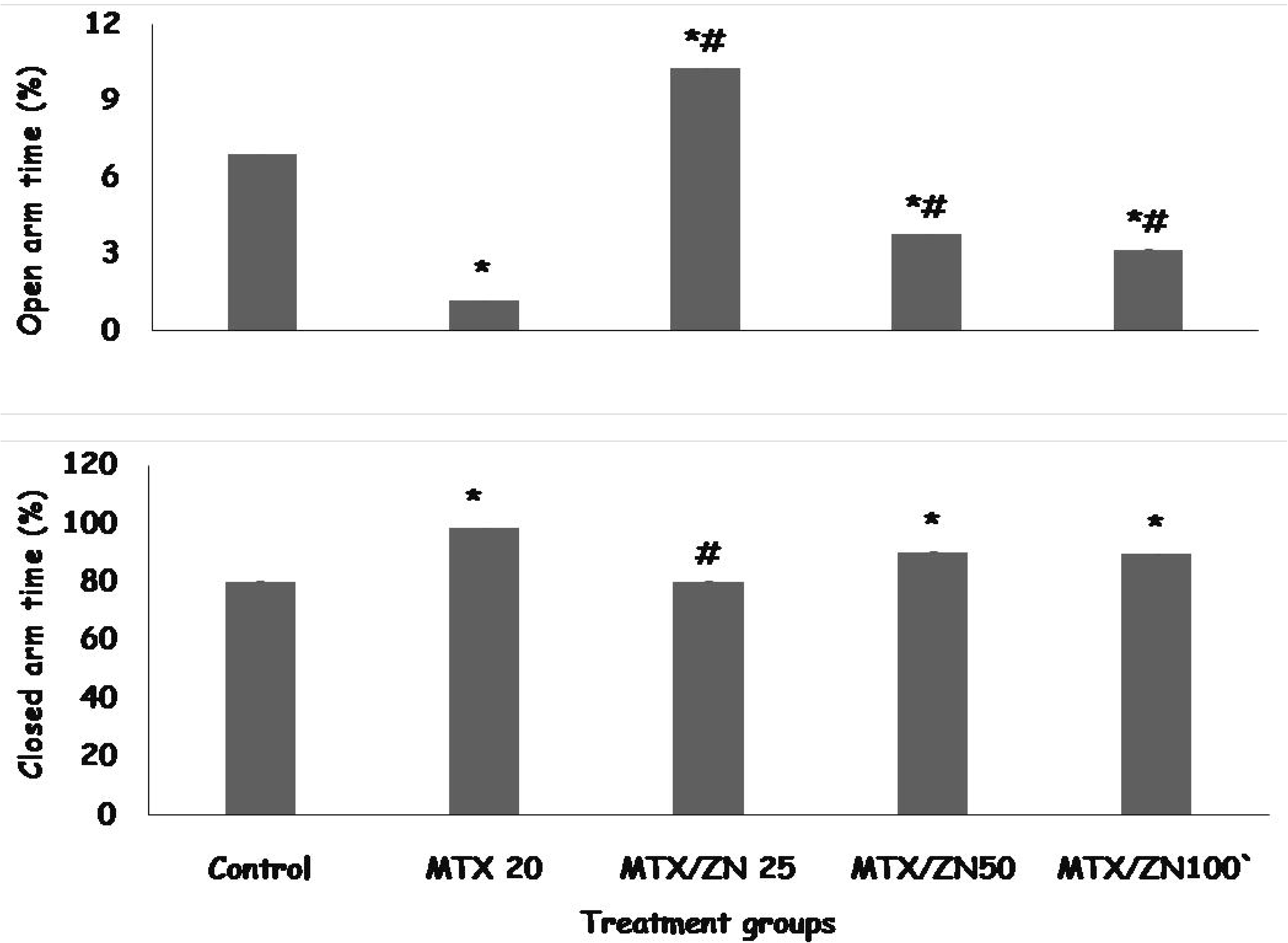
Effect of zinc on Neuron specific enolase immunoreactivity in the Dentate gyrus region of the hippocampus. A: Control, B: Methotrexate Control. C: MTX/ZN25. D: MTX/ZN50, E: MTX/ZN100. Photomicrograph showing small pyramidal cells (sPc), and neuroglia scattered within the molecular layer (ML). MTX: Methotrexate. ZN: Zinc. Neuron specific Enolase immunoreactivity, NRC: Neuron specific enolase reactive cells, Ng: Neuroglia, GC, Granule cells. Magnification=x160

## Discussion

This study examined the protective ffects of dietary zinc supplementation on body weight, food intake, brain function, oxidative stress markers, antioxidant status, inflammatory parameters, hippocampal histomorphology and neurotransmitter levels in adult rats. The results of the study revealed an increase in body weight and food intake, and improvement in antioxidant status; memory scores, neurotransmitters and interleukin levels.

In this study, the supplementation of zinc in methotrexate treated rats was associated with a concentration dependent decrease in body weight and food intake compared to control and a concentration dependent increase compared to methotrexate control. The effects of zinc on body weight and food intake have been reported severally [33, 56–59]. Zinc has been shown to cause either an increase in body weight or a decrease depending on the basal weight of the subject [33, 56–59]. In this study methotrexate therapy resulted in significant weight loss corroborating the results of another study by Moghadam et al. [60] that revealed that the administration of methotrexate was associated with a decrease in body weight. It also decreased food intake consistent with other studies that had reported that methotrexate decreased food intake in addition to altering the gastrointestinal tract integrity [61]. Dietary supplementation with zinc was observed to reverse the weight loss caused by methotrexate. Zinc supplementation also reversed the reduction in food intake that was associated with methotrexate therapy. Zinc’s ability to mitigate the effects of methotrexate on body weight and food intake can be attributed to its effect oxidative stress, inflammation and its ability to modulate lipid and glucose metabolism and regulate insulin expression [62].

The neurotoxic effects of methotrexate has been reported severally [63–68]. Methotrexate has been shown to cause self limiting episodes of seizures, confusion, headaches and focal weakness [69]. Also documented were reports of cognitive deficits including memory impairment and impairments attention and executive functioning [68, 70]. In this study we observed that methotrexate administration was associated with central inhibitory effects as evidenced by a decrease in horizontal locomotion, rearing and grooming. Also observed was a decrease in spatial working memory and increase in depression and anxiety-related behaviours. The effects of methotrexate on the brain have been attributed to its ability to induce oxidative stress, inflammation inhibit neurogenesis and alter neurotransmitter activity [70]. In groups that received dietary zinc supplementation a mitigation of the deleterious effects observed with methotrexate were observed at the lowest dose of zinc while a worsening of the effects observed with methotrexate were observed at the two higher doses, suggesting a dose dependent response with dietary zinc supplementation. The effects of zinc on behavioural indices has been examined severally [63–71]; results from experiments carried out in our laboratory [33, 71, 72] revealed administration of zinc in health [72] or following the induction of disease [33, 71], was associated with dose, concentration or time dependent central inhibitory effects. There was also evidence of reversal of memory impairment and significant anxiolytic behaviour supporting our observations in this study. The effects of dietary zinc supplementation on neurobehaviour can be attributed to its ability to modulate both excitatory and inhibitory brain neurotransmitters [71].

The mechanistic effects of Zinc on the brain has been studied effectively, there are reports that one of the key functions of free zinc within the synaptic vesicles in the brain is the modulation of a number of brain receptors and neurotransmitters. The effects of zinc on the activities of glutamate, gamma amino butyric acid and glycine have been reported [73, 74]. In this study the effects of dietary zinc supplementation on hippocampal levels of acetylcholine, dopamine, serotonin and BDNF in methotrexate treated rats was examined. Methotrexate therapy was associated with a decrease in the hippocampal levels of acetylcholine, dopamine, BDNF and an increase in serotonin levels. This corroborates the results of studies that had reported that methotrexate reduced the concentration of dopamine and serotonin [75, 76]. The administration of zinc reversed the methotrexate effects on the neurotransmitters, with increase levels of acetylcholine and BDNF which could be responsible for the improved spatial working memory scores observed in the radial arm and Y maze paradigms.

The pro-oxidant and proinflammatory effect of methotrexate was confirmed in this study. The results showed an increase in MDA levels, a decrease in TAC and an increase in the levels of inflammatory cytokine (TNF-α and IL-1β) with a decrease IL-10. Several studies have reported the ability of methotrexate to reduce antioxidant capacity and increase oxidative stress [77, 78] in different tissues Cytokines are considered as signaling molecules that are important the modulation of inflammatory and immune responses. Methotrexate has been shown to possess a dose dependent effect of inflammation; while a number of studies have reported antiinflammatory effects of methotrexate [79, 80], a few other studies have reported a proinflammatory effect [81–83]. Also in this study we observed that zinc supplementation was associated with a reversal of methotrexate induced changes in oxidative stress and inflammatory cytokine The antiinflammatory and antioxidant potential of zinc has been reported severally [84, 85].

There is ample evidence in support of the ability of methotrexate to induce neuromorphological injury in humans and rodents [63–67]. In this study we observed histopathological changes in the hippocampus that were suggestive of neuronal injury, these pathological changes had been preceded by evidence of memory loss in both the radial arm and Y maze paradigms. Also observed were the ability of dietary zinc supplementation to protect against the development of methotrexate induced neuronal injury. At higher concentrations of zinc a worsening of the neuronal degeneration is observed particularly in the CA3 region, There we observed loss of sized pyramidal neurons with an increase in the density of degenerating pyramidal cells with darkly staining nuclei. This histopathological effects observed at higher concentrations of zinc supplementation were also observed in the results of behavioural tests s evidenced by an increased central inhibitory response in the open field paradigm and increased anxiety in the elevated plus maze paradigm when zinc was supplemented at 50 and 100 mg/kg of feed. The histopathological changes observed are also buttressed in this study by its effects in reducing neuron specific enolase reactivity in the hippocampus and restoring memory function in the behavioural paradigms.

Neuron specific enolase (NSE) is an acidic protease enzyme that is unique to neurons and neuroendocrine cells. It is involved in the brain’s glycolytic pathway of energy metabolism. It has been reported to be released from neurons especially during periods of injury and is currently considered a sensitive indicator of the severity of nerve cell injury and prognosis [86]. In this study, an increase in NSE reactivity was observed in groups administered methotrexate, suggesting a possible increase in neuronal injury which is consistent with our observations in the H and E stained sections of the dentate gyrus. In the groups fed zinc diet a reversal of the increased immune reactivity was observed. A few studies have reported that the increased serum concentration or increased expression of NSE occurs in neuronal injury and following glial activation and injury [87–89]. This reactivity could be secondary to alterations in mitochondrial metabolism within the neurons that generate reactive oxygen species thereby enhancing neuronal degeneration. Therefore the effect observed with zinc supplementation in this study could be as a result of zinc’s antioxidant capacity as well as its ability to modulate mitochondrial respiration. These view is supported by the results of a study [90] in a mouse model of Alzheimer’s disease, in which dietary zinc supplementation was shown to restore impaired mitochondrial respiration and also increase levels of BDNF (as was observed in this study). The reversal of methotrexate-induced decrease in BDNF levels that was observed in groups fed zinc could also contribute reduction in mitochondrial dysmetabolism [91] and in our opinion a reduction in NSE reactivity. This notion is supported by reports that BDNF also contributes significantly to mitochondrial dynamics and oxidative efficiency [92].

## Conclusion

In conclusion, the results of this study support a possible protective role for dietary zinc supplementation in chemotherapy-induced neurotoxicity. However, more research is required to ascertain its possible use as a neuroprotective agent in methotrexate-induced neurotoxicity

## Author Contribution Statement

A.Y.O and O.J.O conceived and designed research. AYO, OBO and OJO conducted experiments and analyzed data. A.Y.O and O.J.O wrote the manuscript. All authors read and approved the manuscript.

## Ethics Approval and Consent to Participate

All procedures performed on the animals were in accordance with approved protocols of the Faculty of Basic Medical Sciences, Ladoke Akintola University of Technology, Ogbomoso, Oyo State, Nigeria and within the provisions for animal care and use as prescribed by the scientific procedures on living animals, European Council Directive (EU2010/63).

## Human and Animal Rights

No humans were used in this study. All procedures performed on the animals were in accordance with approved protocols of the Ladoke Akintola University of Technology Consent for Publication:

Not Applicable.

## Availability of Data and Materials

The datasets generated during and/or analysed during the current study are available from the corresponding author on reasonable request.

